# A polydiolcitrate-MoS_2_ composite for 3D printing Radio-opaque, Bioresorbable Vascular Scaffolds

**DOI:** 10.1101/2023.10.27.564364

**Authors:** Beata M. Szydlowska, Yonghui Ding, Connor Moore, Zizhen Cai, Carlos G. Torres-Castanedo, Evan Jones, Mark C. Hersam, Cheng Sun, Guillermo A. Ameer

## Abstract

Implantable polymeric biodegradable devices, such as biodegradable vascular stents or scaffolds, cannot be fully visualized using standard X-ray-based techniques, compromising their performance due to malposition after deployment. To address this challenge, we describe composites of methacrylated poly(1,12 dodecamethylene citrate) (mPDC) and MoS2 nanosheets to fabricate novel X-ray visible radiopaque and photocurable liquid polymer-ceramic composite (mPDC-MoS_2_). The composite was used as an ink with micro continuous liquid interface production (μCLIP) to fabricate bioresorbable vascular scaffolds (BVS). Prints exhibited excellent crimping and expansion mechanics without strut failures and, importantly, required X-ray visibility in air and muscle tissue. Notably, MoS_2_ nanosheets displayed physical degradation over time in a PBS environment, indicating the potential for producing bioresorbable devices. mPDC-MoS_2_ is a promising bioresorbable X-ray-visible composite material suitable for 3D printing medical devices, particularly vascular scaffolds or stents, that require non-invasive X-ray-based monitoring techniques for implantation and evaluation. This innovative composite system holds significant promise for the development of biocompatible and highly visible medical implants, potentially enhancing patient outcomes and reducing medical complications.

**TOC:** 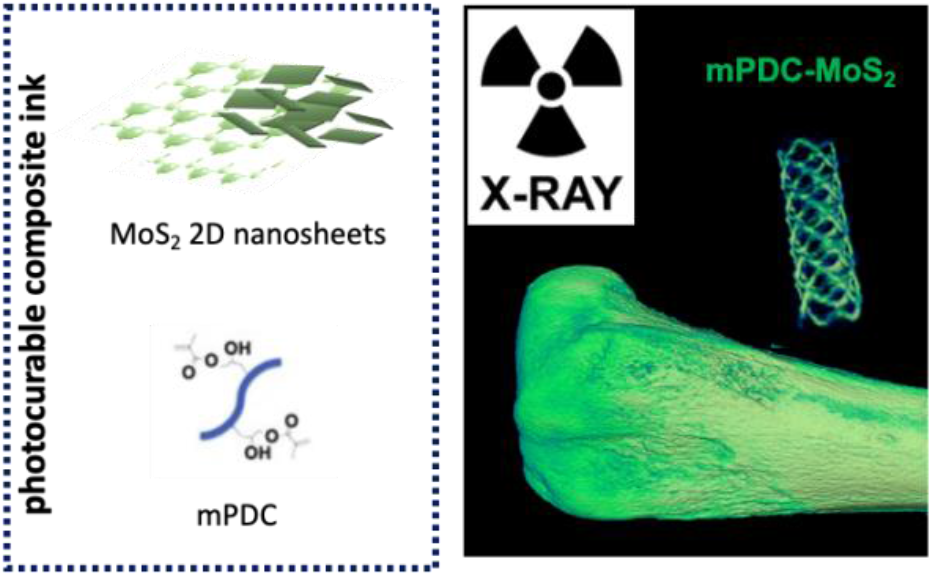

## INTRODUCTION

Arterial occlusions and constrictions, diagnosed as atherosclerotic coronary artery disease and peripheral artery disease resulting from plaque build-up inside blood vessels, impact 274 million people globally.^1^ Current treatment involves placement of a stent – a small tubular grid aiding re-opening and support of the narrowed, occluded vessel lumen with the goal of restoring and maintaining blood flow.^2^ Correct stent placement and expansion are essential to the success of the procedure and long-term outcome. Malposition or inadequate expansion may lead to complications ranging from stent failure to acute thrombosis and an acute myocardial infarction, i.e., heart attack.^2 3^

Considering small dimensions of stents and the target blood vessels in the heart, appropriate placement can be challenging and must be aided by fluoroscopy, an X-ray-based visualization method. Currently, stents with the best fluoroscopic visibility are made of metal in the form of a tubular metal mesh structure.^4^ They provide mechanical support to the vessel but often have long-term complications such as late stent thrombosis with no possibility of future intervention at the same site resulting in impaired vessel biomechanics due to the permanently stented lumen.

To address the aformentioned challenges, interest and research into bioresorbable stents has grown in recent years.^2,5-9^ A bioresorbable stent, referred to also as a bioresorbable vascular scaffold (BVS) provides temporary mechanical support while allowing the tissue to remodel and restore biological function.^10^ However, as they are typically fabricated using polymers (e.g., polylactic acid), they have suboptimal mechanical properties and a lack of intrinsic radiopacity, leading to no or poor X-ray visibility.^10^ X-ray visibility is important for BVS deployment and evaluation post-deployment. Conferring radiopacity to these BVS has been traditionally addressed by attaching heavy metal marker points (e.g., gold or tantalum) to the device.^11^ The procedure is done manually by welding and is both time-consuming and cost-intensive. The position of the BVS can be detected; however, the stent structure remains invisible to the surgeon. Other approaches to increase X-ray visibility of polymeric stents include the incorporation of metallic nanoparticles and more recently described, the clinical contrast agent iodixanol.^11^ The use of metallic nanoparticles results in imaging artifacts, medical complications, or suboptimal stent mechanical properties.^12-14^ Another challenge of BVS is exacerbated inflammation caused by their degradation products.^14^

Recently, we have reported on the fabrication of BVS via micro continuous liquid interface production (uCLIP) using bioresorbable citrate-based biomaterials (CBBs).^15^ In addition to their antioxidant properties and excellent biocompatibility, CBBs have demonstrated compatibility with high-resolution 3D printing processes, facilitating the fabrication of polymeric BVS with individually customized anatomical geometry and size with precision down to 100 μm.^15^ However, X-ray contrast is low and the use of metal markers leads to concerns about residual millimeter-scale metal pieces in the body post-resorption of the polymer matrices. To address this issue, we recently reported the incorporation into the BVS of the clinical contrast agent iodixanol (Visipaque, GE Healthcare), a USA FDA-approved X-ray contrast agent, with minimal impact on required BVS mechanical properities.^12,15^ Although X-ray visibility was obtained, additional research is needed to increase the contrast density in the material and understand the effects of iodixanol on the surrounding tissue to enable clinical use. Therefore, additional materials-based strategies are needed to realize the full potential of fluoroscopically visible BVSs.

Herein, we report the use of MoS_2_ nanosheets, a material with a relatively high atomic number (Z=54),^12^ to provide X-ray opacity and reinforce the mechanical strength of the stent. Exfoliated nanosheets are especially promising due to their high surface-to-volume ratio and liquid phase processability. Thus, they can be densely packed into a polymer solution and act as individual nano X-ray markers. Importantly, MoS_2_ has been reported to be biocompatible^16^ and used in scaffolds for tissue regeneration.^17,18^ We show that mPDC-MoS_2_ composite can be processed using uCLIP to obtain high-resolution printing of mechanically competent BVS. mPDC-MoS_2_ BVS exhibit mechanical properties, significant X-ray contrast, biodegradability, and cytocompatibility.

## RESULTS AND DISCUSSION

### 1. MoS_2_ nanosheets are biodegradable and can be incorporated into mPDC ink for uCLIP processing

Molybdenum (IV) disulfide (MoS_2_) nanosheets used in this work as a ceramic radiopaque filler were prepared via. liquid phase exfoliation.^19-22^ Briefly, bulk MoS_2_ powder was immersed in water-surfactant (SC) solution and probe sonicated to break intralayer van der Waals interactions and isolate nanosheets. Since produced dispersion has been known to have polydisperse characteristics (e.g., containing nanosheets of broad size and thickness distribution)^22-24^, a multistep centrifugation process was employed to isolate dispersions of narrow size distribution and desired nanosheets dimensions.^23^ That was followed by additional purification steps where nanosheets were transferred into deionized water and ethanol to remove possible impurities and surfactant excess. Finally, the material was dried into powder for use to create the composite (see *Methods* section for experimental detail).

MoS_2_ nanosheets with well-defined, sharp edges, were observed in SEM images (**Figure 1A)**. Dispersion narrow size distribution was confirmed with statistical analysis of features in SEM images and is reflected in length, <L> histogram (**Figure 1B**) with average <L>, and <N> of 514 nm and 32, respectively. <L> was evaluated statistically with the help of SEM, while <N> was extracted from the extinction spectra.^24,25^ The spectroscopic signature of MoS_2_ is also presented at the extinction spectra, showing transitions characteristic of multilayer material in the exfoliated form (A and B exciton at 695 and 630 nm, multilayers). **(Figure 1C)** ^21^

**Figure 1.**
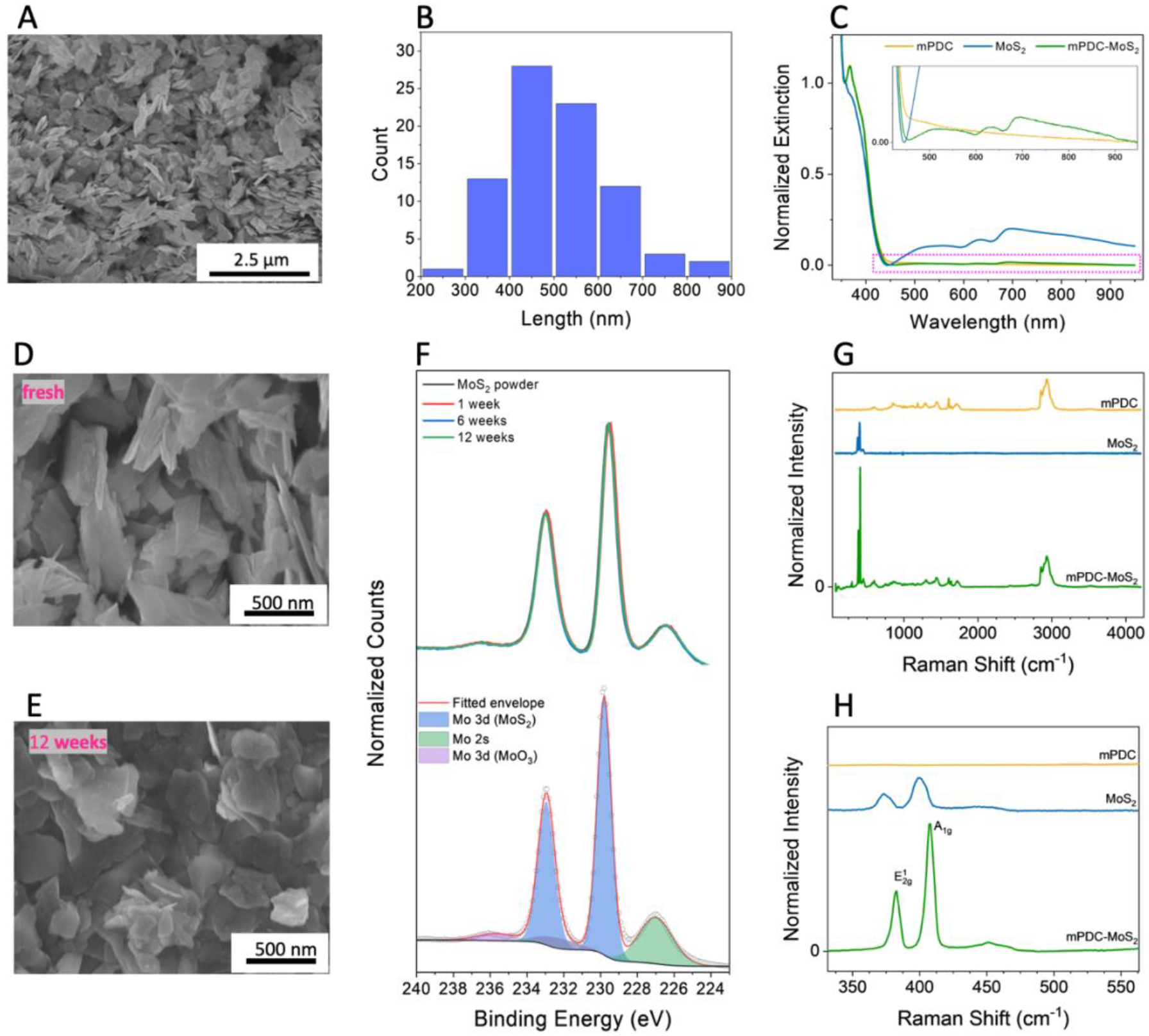
MoS_2_ powder and mPDC-MoS_2_ composite characterization. A) SEM picturing nanosheets of MoS_2_, B) Histogram of nanosheets length distribution in MoS_2_ powder, C) Extinction spectra of mPDC, MoS_2_ and mPDC-MoS_2_, A,D,E) SEM picturing nanosheets of MoS_2_, before and after exposure to 1xPBS at 70°C, F) XPS of MoS_2_ powder, G) Raman Spectra of MoS_2_ powder, printed mPDC, and printed mPDC-MoS_2_, H) Enlarged Raman spectra at low wavenumber range.

It is important for the components of the BVS to be biodegradable over time to minimize chronic foreign body reactions that may lead to reocclusions of the blood vessel. Isolated MoS_2_ nanosheets exhibit a notable level of degradation in the phosphate saline buffer (1x PBS) environment. PBS is commonly used in biological research as it helps to maintain a constant pH, and its osmolarity and ion concentration match those of the human body.^26^ It is visually demonstrated in SEM images (**Figure 1A, D, E top to bottom)** where initially, sharp edges of MoS_2_ nanosheets transform into rounded features as the powder is soaked and stirred in 1xPBS at 70°C (considered accelerated degradation conditions) for an extended period. This observation can be explained as material erosion^27,28^ promoted *via*. elevated temperature and physical agitation.^28^ XPS measurements reveal no chemical changes in the molybdenum (Mo 3d) and sulfur (S 2s) features (**Figure 1F**),^29^ confirms high chemical bio-stability of MoS_2_ nanosheets and degradation mechanism as being of a physical nature only. The lack of chemical degradation eliminates the concern of potential toxic effects of degradation products released into the body. However, the effect of the physical presence of the MoS_2_ in tissue must be fully assessed.

The high quality of the MoS_2_ nanosheets in their pure powder form and post-embedding in the mPDC matrix was confirmed with Raman Spectroscopy (Figure 1G). Raman spectra were obtained from freshly prepared MoS_2_ powder (blue), pure mPDC print (yellow), as well as the MoS_2_-mPDC print test material (green). The first one showcases the characteristic vibrational modes of MoS_2_ (E^1^_2g_ and A^1^_g_), confirming the high quality of the initial material.^30^

Further, spectra corresponding to the MoS_2_-mPDC printed scaffold combine MoS_2_ and mPDC fingerprints (green). Taken together with significantly sharper and more spaced MoS_2_ features, the data confirm the successful dispersion of MoS_2_ in the mPDC matrix, uniformity of the nanophase, and high crystallinity due to post-thermal annealing.^31^ There is a change in the spacing reflecting the elimination of the re-stacked layers present in the initially prepared powder when compared to the composite form (**Figure 1H)**. This change is further confirmed by the extinction spectra of the liquid ethanol-mPDC-MoS_2_ ink (**Figure 1C**). In the high-wavelength range, there is no significant increase in the background, indicating well-dispersed material and material aggregation.^31^ This well-blended ink facilitates the uniform printability of the composite ink, which is desirable for the intended application.

The polymeric ethanol-mPDC-MoS_2_ ink was prepared *via* centrifugal mixing from two main components: mPDC and MoS_2_. mPDC was synthesized as reported in our previous work.^11^ MoS_2_ nanoplatelets were isolated from bulk MoS_2,_ as described in the previous section. The two components were blended at 5 wt% using centrifugal mixing for 5 min with 800 rpms yielding shelf-stable ink. No additional dispersant or stabilizer^32,33^ was used as the prepared ink was stable throughout the printing period.

### 2. mPDC-MoS_2_ composite BVS can be printed via uCLIP

The ink was successfully 3D-printed using the μCLIP process to produce BVSs with strut sizes of 139 μm, as measured with digital optical microscopy and an outer diameter of 2.31 mm and strut thickness of 139 μm based on SEM images (**Figures 2B and 2D**). Following the printing process, stents were heat-treated at 80 °C for 2 hours in argon. Various thermal treatment conditions were explored (see *Supporting Information*, **Figure S1** for details) to assist in residual solvent evaporation and thermal curing *via* a polycondensation reaction.^11^ Similarly, as in our previous work^11^, we observed strut shrinkage (≈10-20%). This effect enhances print resolution, but can be compensated for by increasing the magnitude of the designed dimensions in the CAD model if desired.^11^

**Figure 2.**
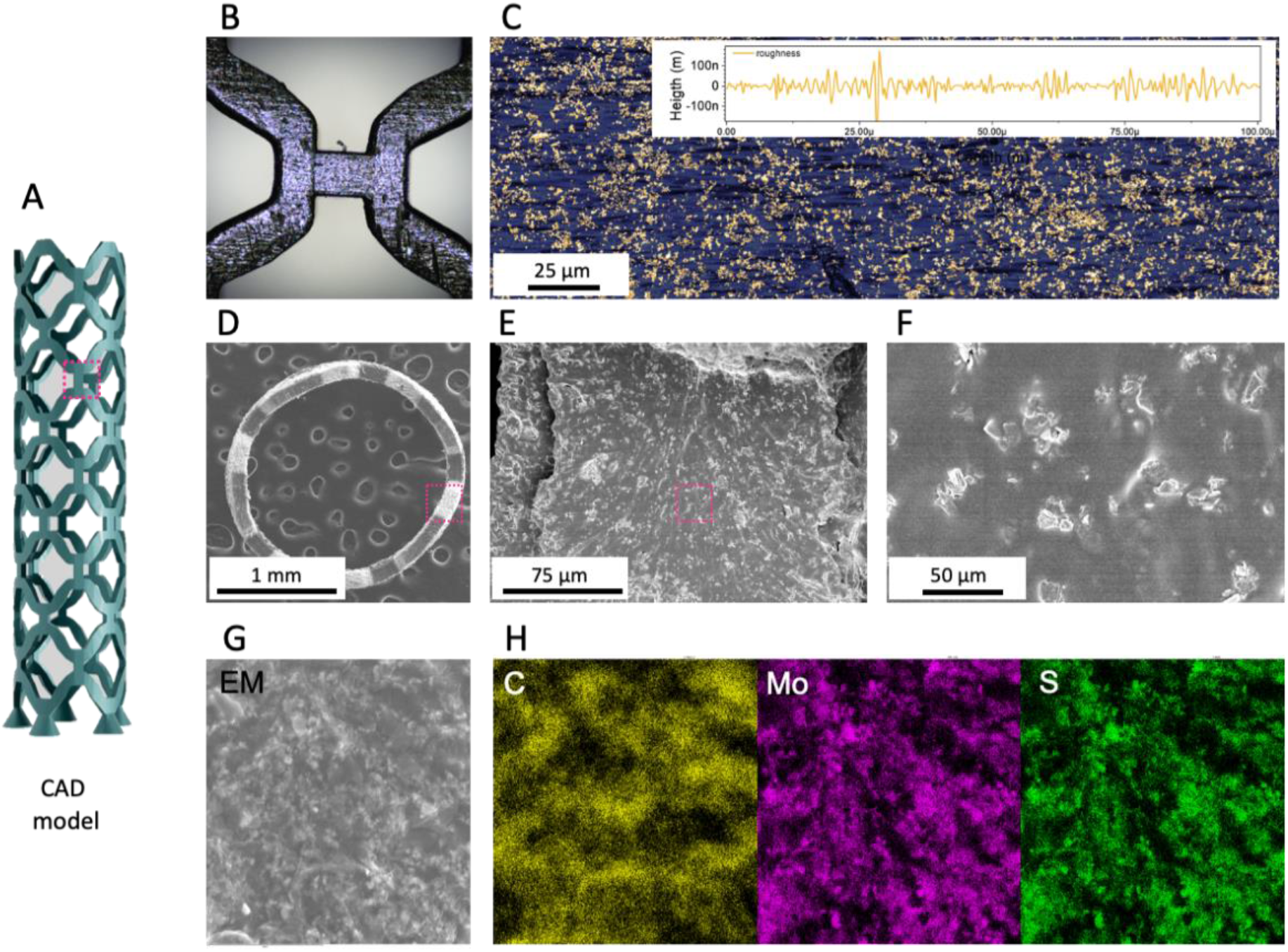
Microscopic Characterization of mPDC-MoS_2_ STENT. A) CAD model of the stents fabricated in this work, with a marked region called *stents knot* (pink frame), B) Optical Microgram of stent’s knot region. C) Height image of the stent surface, inset surface roughness, D, E, F) SEM images of mPDC-MoS_2_ STENT; top view, zoom-ins, G, H) EDS from the mPDC-MoS_2_ STENT surface.

**Figure 2C** presents the height image acquired via confocal microscopy. This figure illustrates a 3D printed MoS_2_-mPDC BVS surface with visible nanomaterial (in yellow) distributed on top of the polymer structure (in navy blue). This mixing results in surface area coverage of 41% and an average print roughness equal to 37 nm, which is higher than the roughness of pure mPDC BVS equal to 2 nm (*see Supporting Information*, **Figure S2**) but still considerably low when compared to the size of the incorporated material (<L> ∼514 nm). These results suggest that most nanosheets are flatly arranged with their basal plane pointing upwards. To gain further insight into the distribution of MoS_2_ within in the printed BVS structures, Scanning Electron Microscopy (SEM) was employed. Cross-sectional images of the BVS are shown in **Figure 2E-F**) These images reveal that MoS_2_ is evenly distributed within the scaffold volume, as is critical to fully harnessing the functionalities added by the presence of MoS_2_. The successful incorporation of MoS_2_ was additionally validated by the presence of Mo (molybdenum) and S (sulfur) signatures in the polymer matrix, as shown in the energy dispersive spectrometry (EDS) mapping and (**Figure 2G-H**). EDS data for the control mPDC stent is described in *Supporting Information* **Figure S3**. Notably, our 3D-printed BVS exhibits no visible stair-casing features as expected from the μCLIP 3D-printing process. This characteristic enables layerless and monolithic fabrication of 3D parts.^11,34^ Consequently, we anticipate isotropic mechanical properties throughout the stent’s geometric structure.

### 3. mPDC-MoS_2_ BVS is Radiopaque

The mPDC-MoS_2_ BVS and pure mPDC BVS (control) were placed in the chicken thigh bone and scanned using micro-computed tomography (CT) to evaluate their radiopacity.^35^ It’s worth noting that these findings pertain to a MoS_2_ loading level of 5 wt%. For results at other loading levels, please refer to *Supporting Information*, **Figure S4**. The resulting computed tomography (CT) image (**Figure 4A**) demonstrates that the MoS_2_-mPDC stent is clearly visible, while the pure mPDC (control) is radio-lucent under the measurement conditions. Moreover, uniform radiopacity along the entire length of the printed device further affirms the printed stent’s homogeneity, particularly in terms of MoS_2_ distribution. This result can be attributed to the high-speed printing process, which effectively suppresses the MoS_2_ nanosheet sedimentation within the ink volume.

Consistent with the substantial visual contrast shown in the radiograph (**Figure 3A**), the quantified relative mean value of Hounsfield units^36^ (in muscle tissue) for MoS_2_-mPDC (5 wt%) BVS is significantly higher than control BVS (mPDC) with values of 1213.4 and -670.1, respectively (**Figure 3B**). In-air radiopacity of the MoS_2_-mPDC (5w%) BVS is equal to 1250 and -650 for mPDC-MoS_2_ and mPDC, respectively. For context, environmental HU values range from -1000 HU for air and 0 for water to ∼2000 HU for dense bone and exceeds 3000 HU for metals.^37^ Noteworthy, commercially available, and FDA-approved commercial agents provide radiopacity at the HU level of -400 and -550 in muscle tissue and in air, respectively. This result places the mPDC-MoS_2_ composite, even with a relatively low (5 wt%) loading of MoS_2,_ in close proximity to the radiopacity expected from the dense bone.

**Figure 3.**
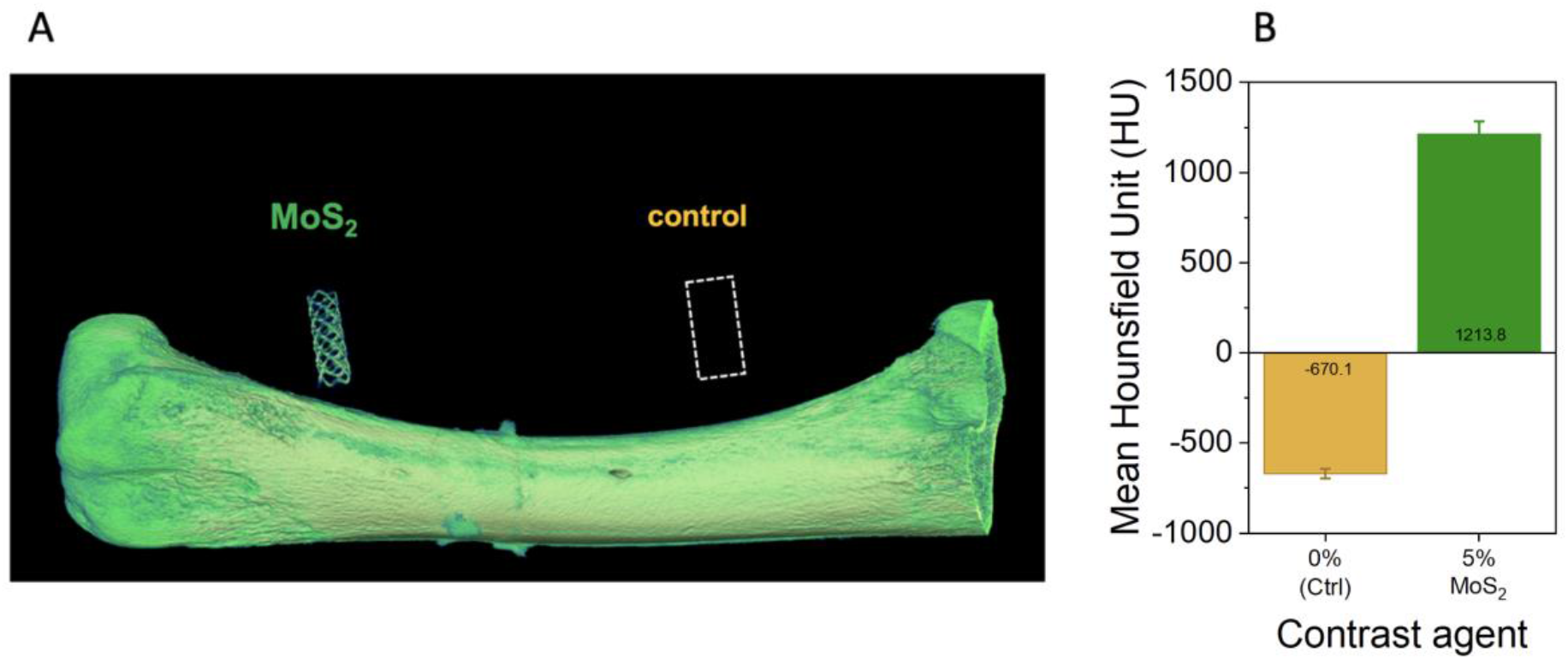
X-ray characterization of mPDC-MoS_2_ STENT via computed tomography (CT). A) CT images (in muscle tissue) of control (mPDC), and MoS_2_ enriched BVS. B) Mean Hounsfield values for the control (mPDC), and MoS_2_ enriched BVS.

Traditionally, individual marker points used for facilitating visualization of the BVS are built from heavy metals such as tantalum or gold. Heavy metals are preferred because they have a high X-ray absorption capacity, resulting in increased mass contrast in imaging.^12,38^ These markers are attached to the stent, providing orientation points for stent positioning.^12^ However, this approach has limitations, as it only visualizes individual points on the stent, making it challenging to visualize the stent’s complete positioning within the vessel. In contrast, the MoS_2_ nanosheets described in this work function as individual nanoscale X-ray markers. Due to their random distribution throughout the mPDC matrix, a consequence of the ink composition and fabrication method – and their high volume-to-mass ratio, these nanosheets allow for the dense packing of MoS_2_. Consequently, the generated X-ray images of the stent demonstrate high fidelity. An additional advantage of using MoS_2_ over metals such as Ta (Z=73) or Au (Z=79) as an X-ray marker in stents is that the MoS_2_ nanosheets (Mo (Z=42) and S (Z=16)) provide X-ray contrast while reducing the propensity of beam hardening (α=Z4λ3)^39^. This effect is prominent in metals and leads to imaging artifacts, necessary algorithmic corrections, and physical filtration in computed tomography.^13,40^

### 4. The MoS_2_-mPDC BVS is biodegradable with required mechanical properties

The mechanical properties of stents play a critical role in their successful and minimally invasive deployment, particularly in supporting surgically treated lumens to prevent collapse. **Figure 5A** shows the original and compressed BVS for both mPDC-MoS_2_ and mPDC (control) material compositions. The radial force was measured by compressing the 3D-printed BVS (within a computer-controlled-diameter cylinder) down to 70% of its original diameter. The results (**Figure 5B**) suggest that all 3D-printed mPDC-MoS_2_ BVS can withstand 30% compression without fracturing. Notably, the radial forces at 30% compression for mPDC-MoS_2_ BVS were comparable to those for mPDC BVS (control), with a slightly higher strength demonstrated by the mPDC-MoS_2_ BVS. The initial radial compression force for dry mPDC-MoS_2_ stents was measured at 2.3 N/mm and decreased to 1.5 N/mm after being soaked in 1X PBS. This value gradually decreased to 0.3 and 0.5 N/mm, indicating the potential for sustained vascular support for at least 140 days. The nominal radial force of mPDC-MoS_2_ BVS at 50% compression, a commonly reported parameter in the literature, was found to be 0.59 N/mm, while mPDC (control) exhibited values of 0.1 N/mm, underscoring the significantly higher strength of MoS_2_-enriched stents (see *Supporting Information*, **Figure S5**).

**Figure 5.**
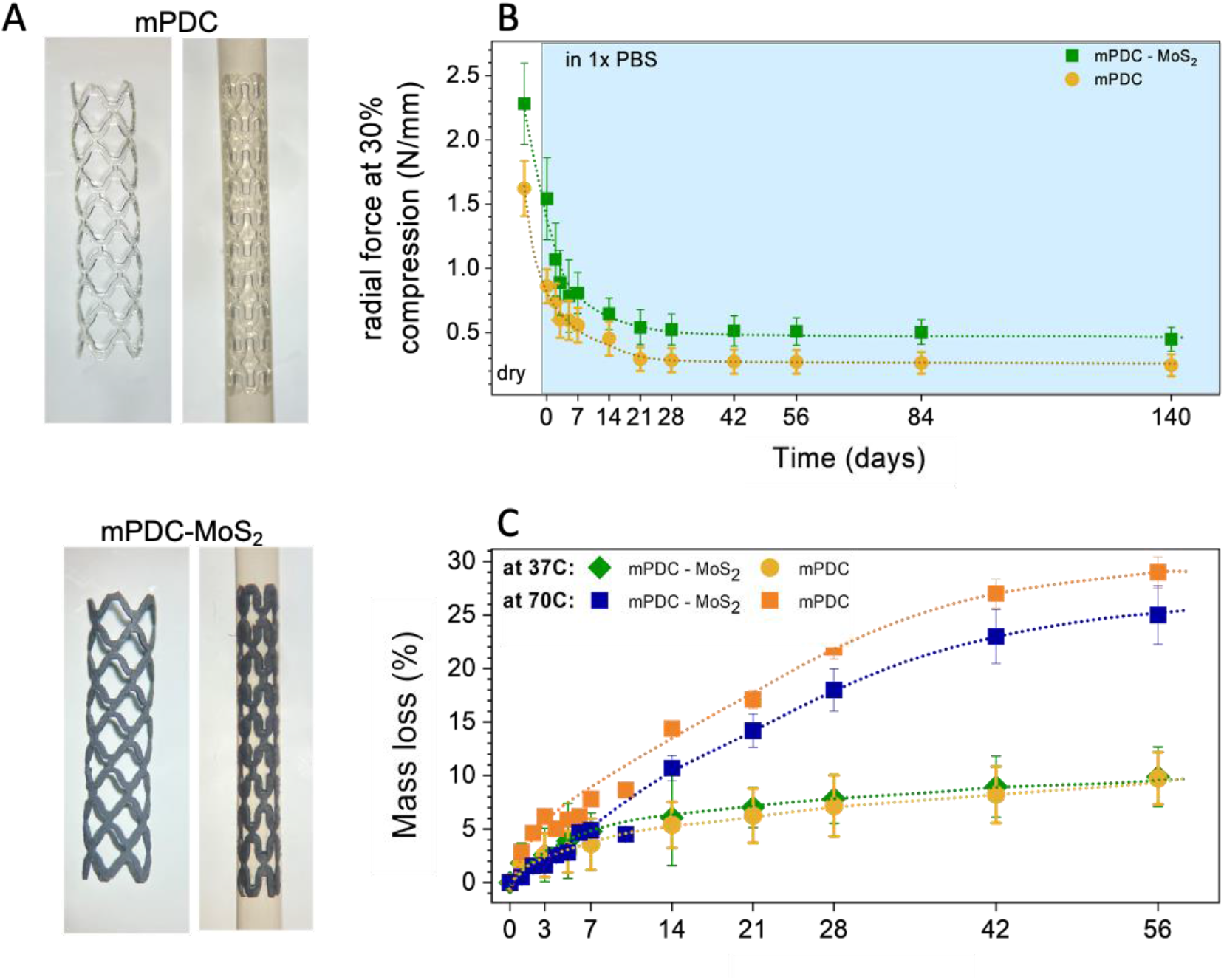
Mechanical Characterization and degradation of mPDC-MoS_2_ and mPDC BVS. A) Optical images of original and crimped mPDC-MoS_2_ and mPDC BVS, B) Radial forces per unit length (N/mm) of mPDC-MoS_2_ and mPDC BVS measured at 30% radial force compression throughout 140 days period (n=9), C) Mass loss recorded mPDC-MoS_2_ and mPDC BVS soaked in 1X PBS at 37C and 70C throughout 56 days period.

Characteristic tensile stress-strain curves of the mPDC-MoS_2_ composite BVS and pure mPDC BVS were acquired using scaled versions of 3D-printed standard dog-bone-shaped samples, which were designed with dimensions proportional to those found in ASTM standard D638.^41^ The MoS_2_-mPDC composite had Young’s modulus of 250 MPa, slightly lower than that of mPDC (280±82 MPa). Simultaneously, the MoS_2_-mPDC composite dog-bones had significantly lower elongation at fracture 12.5% (vs. 28.3±5.0%for mPDC) and ultimate tensile strength of 16 MPa (vs. 10 MPa for mPDC), suggesting that the addition of MoS_2_ reduced the polymer ductility. These parameters are strongly influenced by thermal curing conditions and have the potential to be fine-tuned via changes in post-curing conditions (see *Supporting Information*, **Figure S6**).

Furthermore, mass loss testing was completed on the BVS. To test mass loss, the 3D-printed structure was soaked in 1X PBS at either body temperature (37°C) or elevated temperature (70°C). The 37°C condition was chosen to mimic stent degradation over time in conditions close to the natural body environment. The 70°C condition was chosen to analyze the scaffold’s accelerated physical degradation.^42,43^ At a time point of two months, mass loss was evaluated as 10 % at 37°C and 25% at 70°C. The partial degradation of the stent over this time period suggests the potential for full degradation of the stent over a prolonged period of time, although further testing is necessary in order to confirm this prediction.

### 5. The mPDC-MoS_2_ BVS is biocompatible

The cytocompatibility^16,44,45^ of this composite was evaluated through extract (**Figure 6 A-B)** and direct contact (**Figure 6 C-D)** methods. Initially, mouse fibroblast (L929) cells, which were seeded on polystyrene, were incubated with leached extracts from the 3D-printed mPDC-MoS_2_ and mPDC stents, as well as with pure cell culture medium (**Figure 6 A**). The number of L929 cells was slightly lower in contact with all BVS relative to the pure medium. Hence, L929 cell viability was maintained in contact with the mPDC-MoS_2_ stent. A similar outcome was observed when the cells were directly incubated on the printed mPDC and mPDC-MoS_2_ BVS (**Figure 6 C)**.

**Figure 6.**
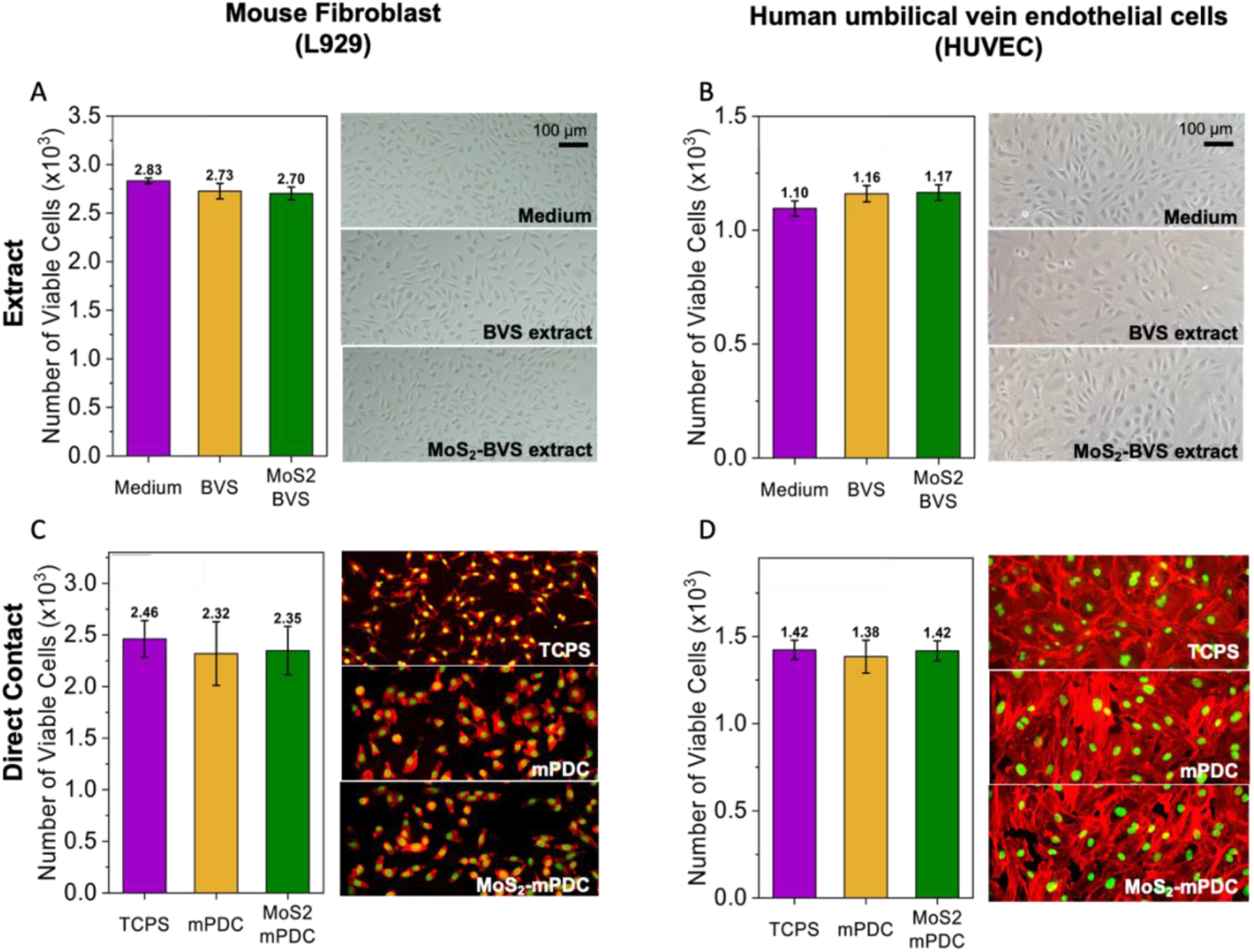
The cytocompatibility of mPDC-MoS_2_ BVS. Mouse fibroblast cells (A, C) and human umbilical vein endothelial cells (B, D) were seeded on tissue culture polystyrene and incubated in (A, B) extract of mPDC-MoS_2_ stents, mPDC stent, control medium, and (C, D) on the top of printed mPDC-MoS_2_ BVS, mPDC BVS, and TCPS.

Similarly, the interaction of mPDC-MoS_2_ BVS with human umbilical vein endothelial cells (HUVEC) was assessed. HUVECs were selected as a test cell type, given their common use in vascular research.^46^ There was no significant difference in HUVEC viability when tested using either method and exhibited typical cobblestone morphology (**Figures 6B and D)**. Therefore, the radiopaque mPDC-MoS_2_ stents showed good cytocompatibility.

## CONCLUSIONS

We describe a novel radiopaque composite material that combines a citrate-based polymer with MoS_2_ nanosheets to achieve strong X-ray visibility while maintaining the required mechanical and cell compatibility properties. The MoS_2_ nanosheets blended into an mPDC matrix at 5 wt% create a promising new material for manufacturing BVSs. The mPDC enables high-resolution 3D printing through light-driven techniques such as stereolithography and confers biodegradability to the printed device. 3D printing enables the use of imaging techniques such as MRI and CT to produce personalized vascular scaffold designs. The enhanced radio-opacity at a low nanomaterial loading of 5 wt% is attributed to the high electron density (Z_Mo_=42) and surface-to-volume ratio of individual nanosheets. SEM analysis confirmed the physical degradation of MoS_2_ nanosheets under physiological conditions, suggesting they may be degraded within the body. Despite the change in nanosheet shapes upon incubation aqueous media, no chemical degradation is observed via XPS, suggesting reduced probability of toxic by-products during scaffold degradation. CT imaging demonstrated excellent X-ray visibility within muscle tissue, which will facilitate BVS placement and long-term monitoring within the human body.

## ACKNOWLEDGEMENTS

**This work was funded in part by grant 5R01HL141933 from the National Heart Lung and Blood Institute. BMS** thanks Deutsche Forschungsgemeinschaft (DFG) for funds within the framework of the Benjamin Walter Fellowship (agreement SZ 463/1-1).The MoS_2_-polymer composite ink development work was supported by the Materials Research Science and Engineering Center of Northwestern University (NSF DMR-2308691). This work was supported in part by the Center for Advanced Regenerative Engineering. The authors thank Caralyn Collins and Eden Taddese for proofreading the manuscript.

## METHODS

### Materials

all chemicals were obtained from Millipore Sigma unless otherwise noted.

### Liquid Phase Exfoliation

MoS_2_ was exfoliated with the assistance of a point probe sonicator. MoS_2_ powder (30g/L) was mixed with Sodium collate serving as a dispersant and stabilizer (8g/L) and dispersed in di-water. The mixture was then sonicated for 2 hrs. Later, the dispersion was centrifuged for 1hr at 6000g, and the supernatant was discarded. Sediment was redispersed in sodium cholate (2g/L) and further sonicated for 6 hrs.

### Size Isolation

Dispersion yielded by Liquid Phase Exfoliation characterized by broad size distribution was run through a centrifugation step to isolate desired nanosheets.

Centrifugation at 100g was performed for 2.5 hrs., sediment was discarded, and supernatant was again centrifuged at 500g. This time, sediment was collected as a desired nanomaterial to form the composite.

### Purification

Nanomaterial collected in the size selection process was further purified by iterative solvent transfer. This was performed twice with DI water and twice with ethanol. The final dispersion of MoS_2_-ethanol was left at room temperature to form a powder MoS_2_ sample.

### Polymer (mPDC) synthesis

Polymer preparation: Methacrylated poly (1,12 dodecamethylene citrate) (mPDC) was synthesized by following the same protocol as reported in previous studies. Briefly, citric acid and 1,12-dodecanediol were melted (165 ÆC, 22 min) in a 2:1 molar ratio, co-polymerized (140 ÆC, 60 min), purified, and freeze-dried to yield PDC pre-polymer. Every 22 g PDC pre-polymer was dissolved in tetrahydrofuran (180 mL) with imidazole (816 mg) and glycidyl methacrylate (17.8mL), reacted (60 ÆC, 6 h), purified, and freeze-dried to yield mPDC pre-polymer.

### Ink Formulation: mPDC ink (control)

75 wt.% mPDC was mixed with 2.2 wt.% Irgacure 819, acting as a primary photoinitiator, and 3.0 wt.% Ethyl 4-dimethylamino benzoate (EDAB), acting as a co-photoinitiated, in a solvent of pure ethanol. **mPDC-MoS**_**2**_ **composite ink**, 5 wt.% of MoS_2,_ and 1 wt.% stearic acid were mixed with prepared PDC control ink with the aid of a centrifugal mixer.

### SEM

Samples were first sputter-coated with gold and platinum before SEM and EDS examination. The structure and morphology of the 3D-printed stent were then examined using SEM (Hitachi SU8030, Japan). The SEM images were analysed using ImageJ^®^ (National Institutes of Health) to measure the strut thickness and diameter of stents. Powder MoS_2_ nanosheets were drop casted onto copper tape prior to imaging. No additional coating was required for the sample preparation.

### EDS

elemental analysis and energy-dispersive X-ray spectroscopy (EDS). The elemental analysis was performed by an EDS detector (Oxford AZtech X-max 80 SDD).

### UV-Vis Spectroscopy

**O**ptical extinction spectra were measured with a UV–-vis spectrophotometer (Cary 5000, Agilent) using quartz cuvettes (path length = 0.4 cm).

### Raman Spectroscopy

Raman spectra were obtained using a Horiba XploRa PLUS microscope (Horiba, Kyoto, Japan) with a 532 nm laser and 1800 mm^-1^ grating. Each spectrum was obtained with 10% laser power and averaged over five acquisitions, each with a duration of 10 s.

### X-Ray Diffraction Spectroscopy

the spectra were collected in a Thermo Scientific ESCALAB 250Xi XPS spectrometer equipped with a monochromatic Al Kα X-ray source (1486.6 eV). The analysis was performed using the Avantage (Thermo Scientific) software. All the peaks were charged-corrected to adventitious carbon (C 1s) at 284.8 eV.

### Thermal Annealing

**T**hermal annealing was performed in a home-built tube furnace with an argon atmosphere. The stents were placed in the tube and vented with pure argon for 30 minutes. The temperature was then ramped to 80C in 20 mins, held for 2 hrs., and finally cooled to room temperature.

### Optical Confocal Microscopy

An Olympus 3D Laser Confocal Microscope was used to evaluate print thickness and surface roughness in a non-contact manner. Images of 50 x 50 μm were taken in a 3D mapping mode and later post-processed with ImageJ^®^ software to extract the thickness and roughness value.

### Computed Tomography (CT)

Measurement of radiopacity by Micro-Computed Tomography (CT) Imaging:

Samples were scanned in a micro-CT scanner (nanoScan PET/CT, Mediso-USA, Boston, MA). Data was acquired under the following parameters: X-ray tube voltage of 50 kVp, 1Å∼1 binning, 720 projection views over a full circle, and 300 ms exposure time. The projection data was reconstructed with a voxel size of 34 μm (in all directions) and using filtered (Butterworth filter) back-projection software from Mediso. Amira 2021.2 (FEI Co, Hilsboro, OR) was used to segment the stents, followed by 3D rendering. The radiopacity of the specimens was quantified using the mean intensity of segmented stents in Hounsfield Units for statistical analysis.

### Mechanical Properties: Young’s modulus and elongation at fracture

The dog-bone-shaped samples were prepared with the same 3D printing and post-processing conditions as the BVS. A tensile test was performed using an Instron universal testing machine (Model 5940, Instron, High Wycombe, UK) equipped with a 2 kN load cell at a crosshead displacement speed of 1 mm/min, which conformed to ISO 527:2012. **Radial force at 50 % compression:** A stent or BVS was placed between two parallel plates of the Instron universal testing machine (Model 5940, Instron), and its resistance force was measured while the sample was compressed down to 50% of its original diameter at a crosshead displacement speed of 1 mm/min (conformed to ISO 25539-2). The radial force was measured and normalized by the respective stent or BVS length and given in N/mm. **<u>Radial force at 30% compression:</u>**

### Mass Loss

Mass loss was assessed by tracking BVS weight at various time stamps, e.g., after 1 day, 2 days, 3 days, 5 days, 1 week, 2 weeks, etc. Stents were soaked in 1xPBS and kept at 70 °C. At the time of mass measurement, they were removed, soaked off, and dried for 5 hrs. prior to mass readout.

### Cytotoxicity Test

**H**uman umbilical vein endothelial cells (HUVECs; ATCC, Manassas, VA) were expanded in growth media consisting of Endothelial Cell Growth Kit-VEGF (ATCC PCS-100-041) in Vascular Cell Basal Medium (ATCC PCS-100-030) under the standard culture condition (37 ÆC with 5% CO_2_ in a humid environment) to 80% confluency before passaging. HUVECs at passages 5–7 were used.

