## Supporting Information for "A polydiolcitrate-MoS_2_ composite for 3D printing Radio-opaque, Bioresorbable Vascular Scaffolds"

Stent, Bioresorbable, $MoS_2$, 2D material, X-ray contrast, Radio-opacity, bio-composite,

### 1. Thermal treatment

mPDC-MoS2 stent prints have been annealed in several conditions with varied times, temperatures, and atmospheres. An example of the influence of the thermal treatment atmosphere on the chemical composition of the final product (stent) is visualized with XPS below. In this case, in air (presence of oxygen) and in argon (no oxygen).

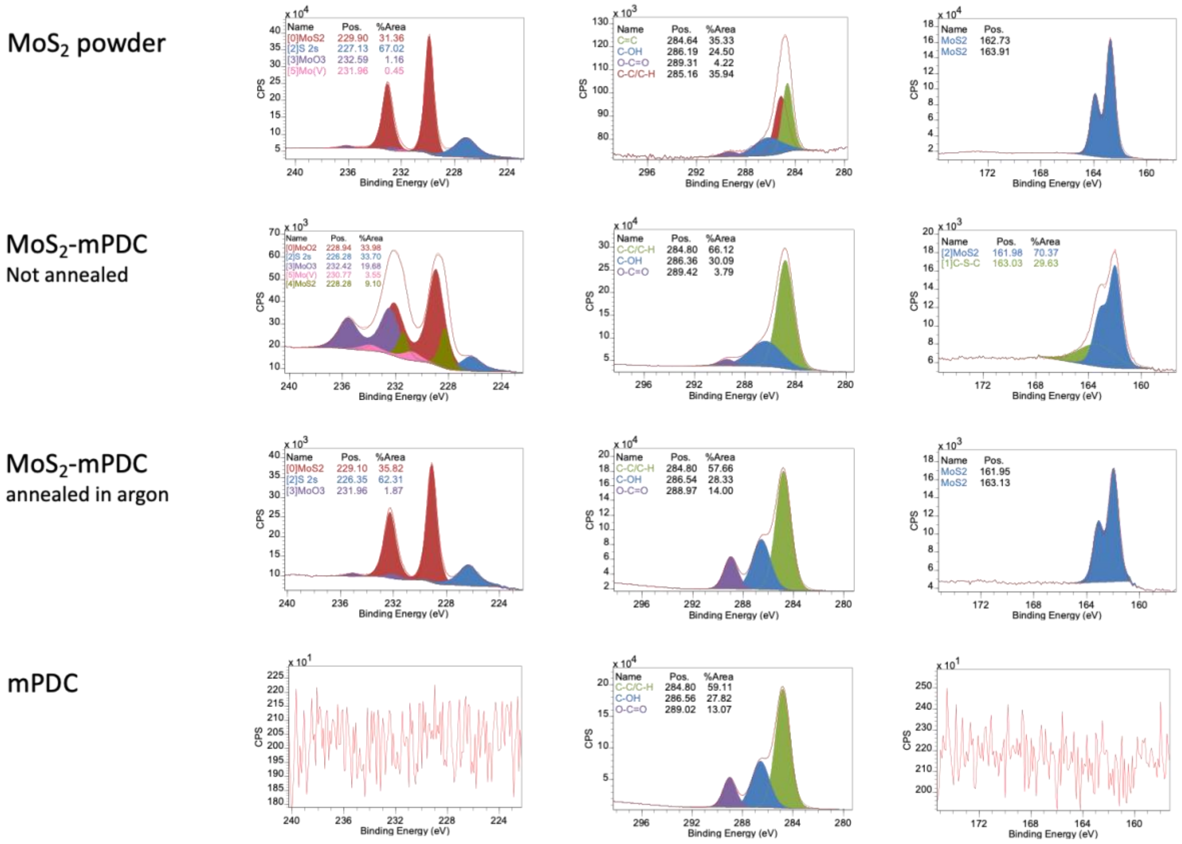


**Figure S1**. **XPS of Mo, S, and O** captured from: 1st row) $MoS_2$ powder, 2nd row) $MoS_2$-mPDC annealed in air, 3rd row) $MoS_2$-mPDC annealed in argon, 4th row) mPDC.

2. **Print Surface morphology**

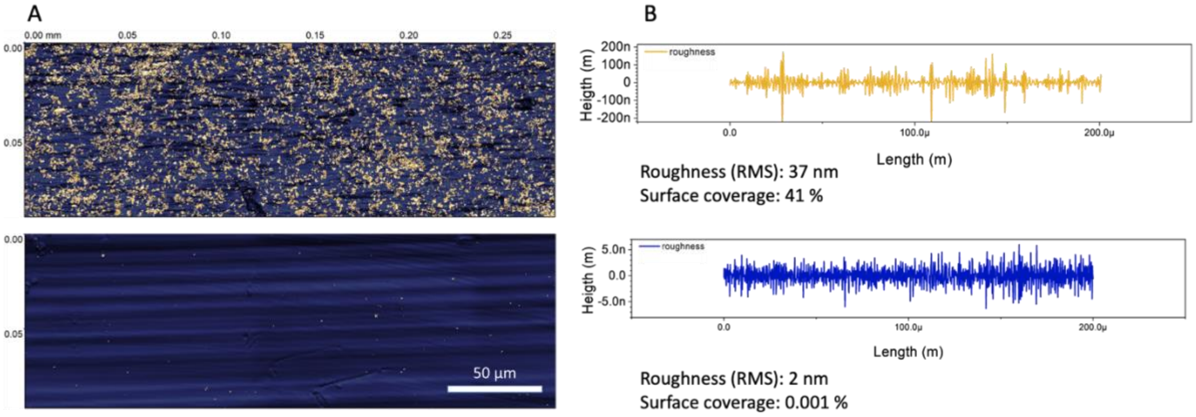


**Figure S2. Characterization of mPDC-$MoS_2$ STENT surface morphology.** A) Height microgram of STENT surface: mPDC-$MoS_2$ (top) and mPDC (bottom). B) Averaged roughness profile of STENT surface: mPDC-$MoS_2$ (top) and mPDC (bottom).

3. **Microscopic Characterization of mPDC STENT (reference sample).**

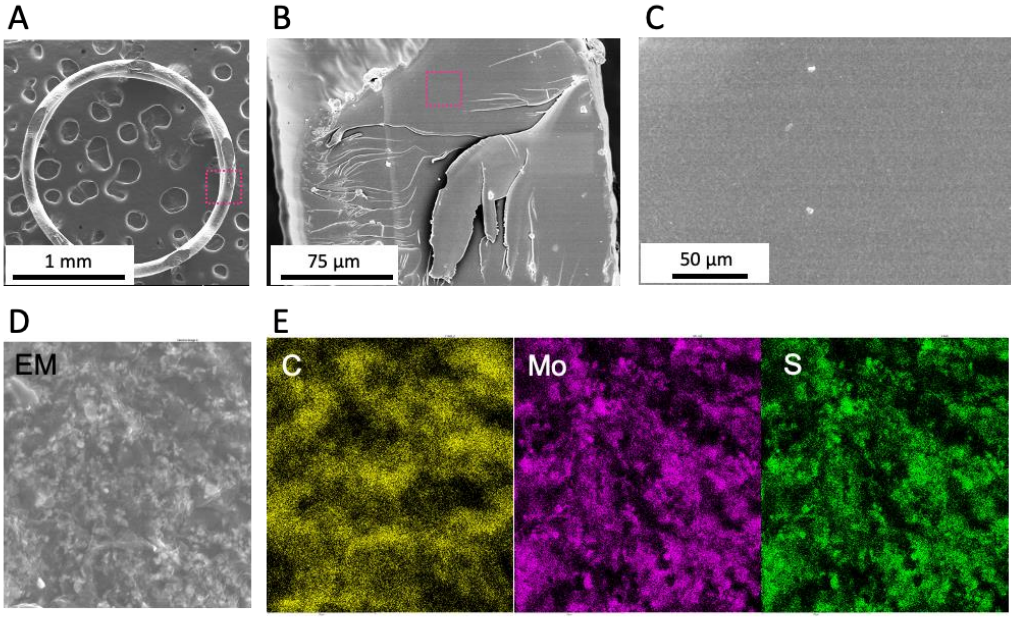


**Figure S3. Microscopic Characterization of mPDC STENT (reference sample).** SEM images of mPDC-$MoS_2$ STENT; top view, zoom ins, F,G) EDS from the mPDC-$MoS_2$ STENT surface.

4. **Radio-opacity – loading level**

Computed Tomography image of stents printed of mPDC-$MoS_2$ composite at different $MoS_2$ loading levels, e.g., 0, 2.5, 5 w%, and 5 w% Iodixanol.

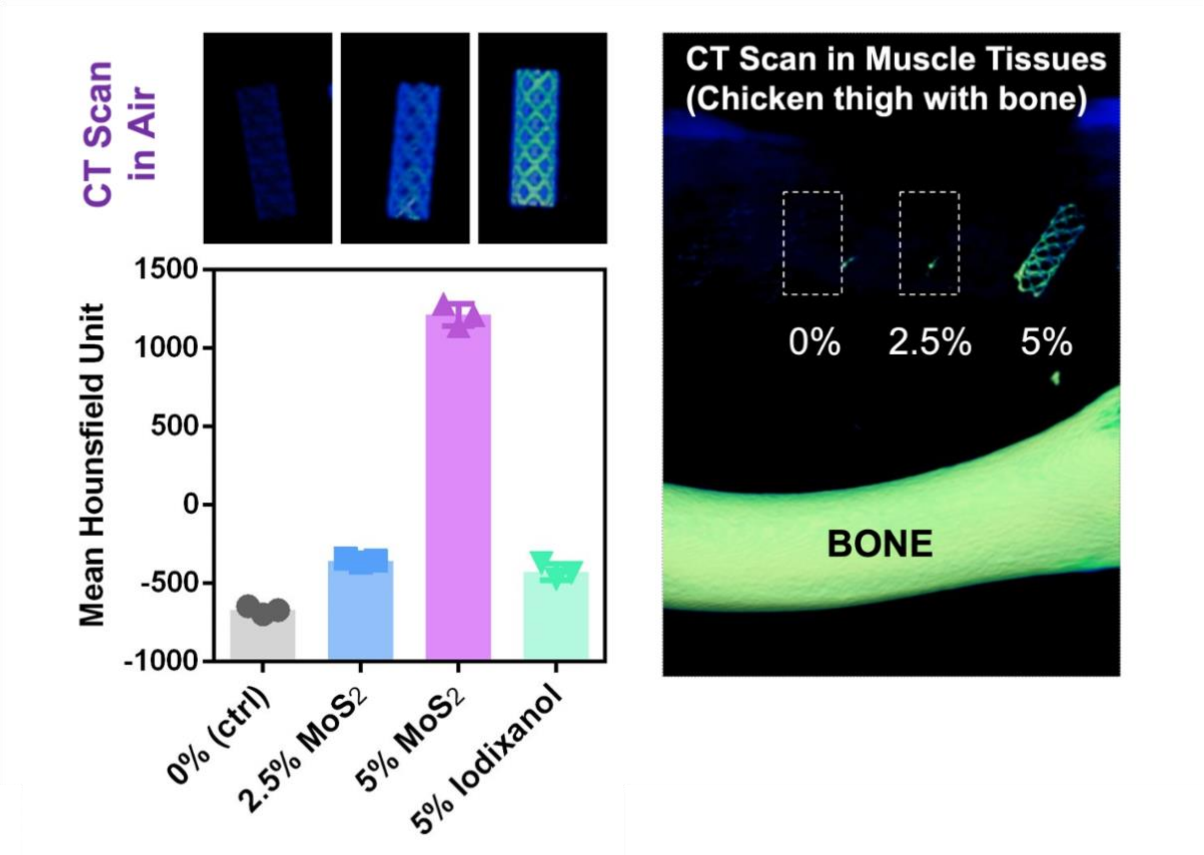


**Figure S4. Radio-opacity evaluation via. Computed Tomography**. Left top ) CT image in the air for 0, 2.5, and 5 w% loading level of $MoS_2$ composite in the printed stent, Left bottom) HU values for 0,2.5, 5 w% loading level of $MoS_2$ composite in the printed stent as well as for 5w% loading of Iodixanol, right) CT image of the same 0, 2.5, and 5 w% loading level of $MoS_2$ composite in the printed stent in muscle tissue.

### 5. Radial compression 50% - the impact of nanomaterial lateral size

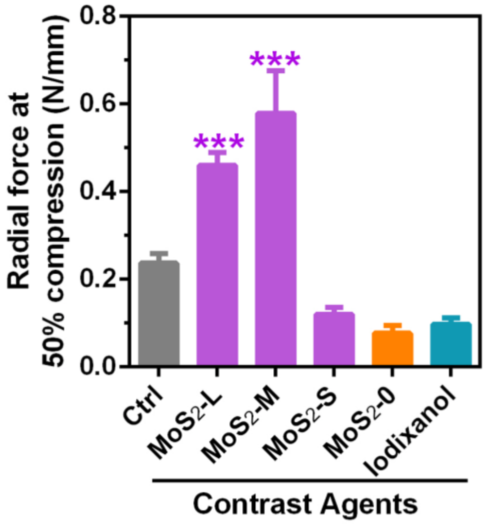


**Figure S5.** Radial forces per unit length (N/mm) of mPDC-$MoS_2$ loaded with 5w% of $MoS_2$ of different lateral size, mPDC and mPDC-Iodixanol stent measured at 50% radial force compression (n=5)

### 6. Radial compression 50% - the impact of the thermal treatment

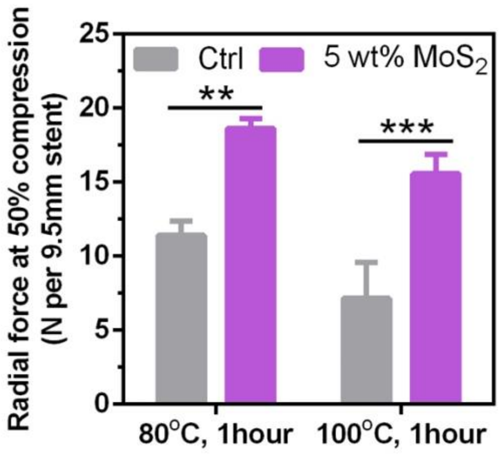


**Figure S6.** Radial forces per unit length (N/mm) of mPDC-$MoS_2$ loaded with 5w% of $MoS_2$ and mPDC stent post-thermal treatment at different temperatures measured at 50% radial force compression (n=5)
